# Habitat preference influences response to changing agricultural landscapes in two long-horned bees

**DOI:** 10.1101/2022.04.15.488450

**Authors:** Gaku S. Hirayama, Atushi Ushimaru

## Abstract

Agricultural intensification and urban development have drastically influenced pollinators living in semi-natural grasslands. Pollinators are likely to have different responses to these land-use changes; some decline rapidly while others maintain or increase their populations. We predicted differences in interspecific response to land-use changes are partly attributed to differences in habitat preference. We examined the distribution and flower use patterns of two closely related *Eucera* species with different habitat preferences. Study sites were meadows surrounding traditional, intensified, and urbanised agricultural lands in the Osaka-Kobe metropolitan area, Japan. Forest-associated *Eucera nipponensis* were significantly fewer in consolidated and urbanised meadows than traditional meadows, whereas open land-associated *E. spurcatipes* exhibited the opposite. A significant negative relationship between their abundances was also found. Both species foraged on legume flowers most frequently (83.5%), but floral use pattern was significantly different in traditional and consolidated meadows. Bee species preferring stable habitats were vulnerable to land consolidation and urbanisation, while species associated with disturbed habitats maintained or increased population size in meadows with the land-use changes. Thus, recent land uses may have different impacts on species with different habitat preferences.

## Introduction

Global pollinator declines caused by land-use changes, such as agricultural intensification and urban development, have become prominent in semi-natural and natural ecosystems (Kearns et al. 1998; Potts et al. 2010; Harrison and Winfree 2015). Traditional semi-natural grasslands surrounding agricultural lands are known to maintain diverse pollinators by providing rich floral resources and nesting sites (Ockinger and Smith 2007). However, recent land-use changes have largely reduced the quantity and quality of such semi-natural habitats, causing declines in wild pollinators, such as bees and syrphid flies (Biesmeijer et al. 2006; Winfree et al. 2009). Although many studies have examined the effects of agricultural intensification or urbanisation on pollinators of semi-natural grasslands separately (Hernandez et al. 2009; Matteson et al. 2008; Martins et al. 2017; Shi et al. 2021), few have examined their combined effects (Olivier et al. 2020).

Agricultural intensification and urban development typically diminish populations of diverse pollinator species (Ollerton et al. 2014; Harrison and Winfree 2015), however, some species maintain or even increase their abundance under such changes (Cane et al. 2008; Matteson et al. 2008; Potts et al. 2010; Martins et al. 2017). Understanding which traits are important in determining responses to the intensified land uses is critical to pick out conservation priority species in land-use changed areas. Interspecific differences in susceptibility to these land-use changes may partly depend on differing habitat preferences among pollinator species (Winfree et al. 2011). For example, it is predicted that species inhabiting relatively stable, natural or semi-natural habitats would be negatively influenced by land-use changes, whereas those preferring more disturbed habitats would become dominant in human-modified landscapes. This prediction is ideally examined by comparing closely related species with similar ecological and morphological traits but different habitat preferences.

To test this prediction, we compared the distribution of two closely related long-horned bee species, *Eucera nipponensis* (Pérez) and *E. spurcatipes* (Pérez), which often coexist in semi-natural grasslands in rural areas (Fig. S1a,b). *E. nipponensis* is typically distributed throughout natural or semi-natural forests and nests on forest floors (Inoue et al. 1990; Hattori et al. 2015; Sugiura and Maehara 2019), while *E. spurcatipes* is exclusively observed in open landscapes, including coastal and residential areas, which are more frequently disturbed (Saito et al. 1991; Hisamatsu et al. 2011; Ogawa and Miyake 2020). Both species are ground-nesting, solitary bees that are widely distributed throughout Japan and have a very long proboscis and a similar foraging period (April to July) (Mitai 2014). Thus, the two species exhibited an overlap in morphological traits and life cycles but differ in habitat preference. Here, we expected that *E. nipponensis* would be more vulnerable to agricultural intensification and urban development than *E. spurcatipes*.

To examine the effects of land-use changes on bee distribution, we surveyed the abundance and flower use of the two *Eucera* species in semi-natural grasslands (meadows) on the levees of traditional, land-intensified (consolidated), suburban, and urban paddy fields. We also investigated the flowering plant richness and abundance and surrounding landscapes for each grassland to examine the factors influencing the distribution of each species. We addressed the following questions: (i) Does *E. nipponensis* decrease in abundance in consolidated and urbanised grasslands? (ii) Does agricultural intensification and urbanisation have any positive effect on *E. spurcatipes* abundance? (iii) Are the impacts of land-use changes caused by changes in surrounding landscapes or floral resources?

## Materials and Methods

### Study area, paddy fields, and plots

The study was conducted in 28 paddy sites across the rural-urban landscape of the Osaka-Kobe metropolitan area (34°66′–96′ N, 134°94′–135°48′ E; Kobe, Akashi, Sanda, Takarazuka, Nishinomiya, Itami, and Kawanishi cities, Hyogo prefecture; Toyonaka city and Nose town, Osaka prefecture) (Fig. S2). In this study site, consolidated and unconsolidated traditional paddy fields, as well as neighbouring secondary forests, are widely distributed in its rural zones (Uematsu et al. 2010), however, paddy fields and secondary forests have been rapidly decreasing in its urban zones since the 1980s (Tsuji et al. 2011; Uchida et al. 2018). In 2021, we investigated abundance and flower use of the two *Eucera* species in meadows which have been maintained on the levees of paddy fields and irrigation ponds, and at the edges of between paddy fields and secondary forest by periodic mowing in the study sites (Uematsu et al., 2010; Uchida et al., 2018).

The meadows were categorised into four land-use types: traditional, consolidated, suburban, and urban (Fig. S3). Land consolidation converts small, irregular, and poorly drained paddy fields into large, quadrangular, and well-drained fields to improve productivity (Uematsu et al. 2010). We defined suburban and urban categories based on the percentage of surrounding developed land area within 1 km of the centre of each study site (suburban < 60%; urban 60% ≤). In the study area, land consolidation and urban development in the surrounding landscape have caused the loss of plant and butterfly diversity in paddy meadows (Uchida and Ushimaru 2014; Uchida et al. 2018). At each study site, we established a 2 × 500 m² belt plot on meadows. We examined a total of 28 plots; seven sites for each of the four land-use types.

### Landscape variables around study sites

For each study site, we calculated the total areas of traditional, consolidated and abandoned paddy fields; other farmlands; secondary forests; water surfaces (rivers and ponds); and developed lands (residential/partly commercial, industrial land uses, and areas for public facilities such as stations, city halls, and hospitals) within a 1 km radius from the site centre using QGIS software (version 3.16.1; QGIS Development Team 2021) and aerial images from Google Maps (2021). We conducted a principal component analysis (PCA) on these seven area variables to reduce the number of landscape variables included in the data analysis. We found that two primary axes explained 63.2% of the total variance (PC1, 46.8%; PC2, 16.4%). The PC1 value was positively correlated with developed lands, water surfaces, and other farmland areas and negatively correlated with forests and consolidated and abandoned paddy field areas. Thus, the PC1 axis indicates changes in the surrounding landscapes from rural with secondary forests and paddy fields to urban with developed lands and farmlands (Fig. S4). The PC2 axis increased mainly with traditional and abandoned field areas and decreased with other farmland areas, indicating the amount of surrounding unconsolidated paddy fields (Fig. S4).

### Bee abundance surveys

We conducted a single bee abundance survey per day for each study site on sunny, warm days from mid-April to mid-May 2021. We observed bees for two hours: one hour in the morning (between 09:00–12:00) and the afternoon (between 12:00– 15:00). We walked along each belt plot, captured all *Eucera* bees using a sweep net (42 cm diameter), and stored them in plastic vials. We held all captured individuals at 1– 4 °C for approximately 10 min in a cooler box. Then, we identified species and made markings with different colours for each species on the thorax of each chilled individual to avoid double counting of the same individuals and released them in the fields (Fig. S1c). All chilled bees recovered well enough to visit flowers within a few minutes. We counted the total number of marked individuals for each species as a measure of the species abundance at each site.

### Flower use surveys

To examine interspecific differences in flower use, we first recorded background flowering entomophilous species and their flower numbers in each plot. Next, we recorded the flower species visited by marked *Eucera* bees and their visit frequency to each species on the same day of the abundance survey. Only visits where bees inserted their proboscis into the corolla tubes or spurs were recorded. We measured the corolla tube length, petal, and inflorescence size (in mm) of four (or all if there were less than four) individuals per flowering species using digital callipers. We followed the method described by Hiraiwa and Ushimaru (2017) to calculate the mean corolla tube length and the visual (attention) size of each species. Flower abundance for each species was calculated as the total visual size of flowers (the mean visual size multiplied by the number of flowers) for each site.

### Data analysis

First, we compared the abundance of the two *Eucera* species among the four land-use types using a generalised linear model (GLM, Poisson errors, and log link function). In the GLM, the land-use type (consolidated; traditional; suburban; urban), bee species identity (*E. nipponensis*; *E. spurcatipes*), and the interaction between them were incorporated as explanatory variables, whereas the abundance of each species for each site was the response variable. To consider spatial autocorrelation in bee abundance, the spatial autocovariates for all sites were calculated using their latitude and longitude measurements and were incorporated as a covariate in the model (Dormann et al. 2007). Second, we examined the effects of landscape variables identified by the PCA analysis (PC1 and PC2) on the abundance of each *Eucera* species using a GLM (Poisson error and log-link function). In this GLM, we included the PC1 value, species identity, their interaction, and the spatial autocovariates as explanatory variables, whereas the abundance of each species for each site was the response variable. We excluded the PC2 value and the interaction between PC2 and species identity in this model because our preliminary analysis revealed no significant effects on bee abundance. Third, we examined the potential of competitive interactions between the two species by analysing a GLM (Poisson error and log link function) in which the abundances of *E. nipponensis* and *E. spurcatipes* were the response and explanatory variables, respectively. In the GLM, we also included the spatial autocovariates of *E. nipponensis* abundance as a covariate. Finally, we examined differences in flower use between the two *Eucera* species for each land-use type using Fisher’s exact test. All statistical analyses were performed using R software (version 3.6.1; R Development Core Team 2019).

## Results

### Bee abundance

We recorded 179 *E. nipponensis* and 373 *E. spurcatipes* individuals at the study sites. *E. nipponensis* was significantly more abundant than *E. spurcatipes* in traditional meadows, whereas this pattern was reversed for other land-use types (Fig. 1; Table S1). PC1 had significant negative effects on the abundance of both species, and the interaction between PC1 and species identity had a significant effect, indicating that *E. nipponensis* decreased more drastically along PC1 gradients (Fig. 2; Table S1). Furthermore, the abundance of the two *Eucera* bees was significantly negatively correlated (Fig. 3; Table S1).

**Fig. 1.**
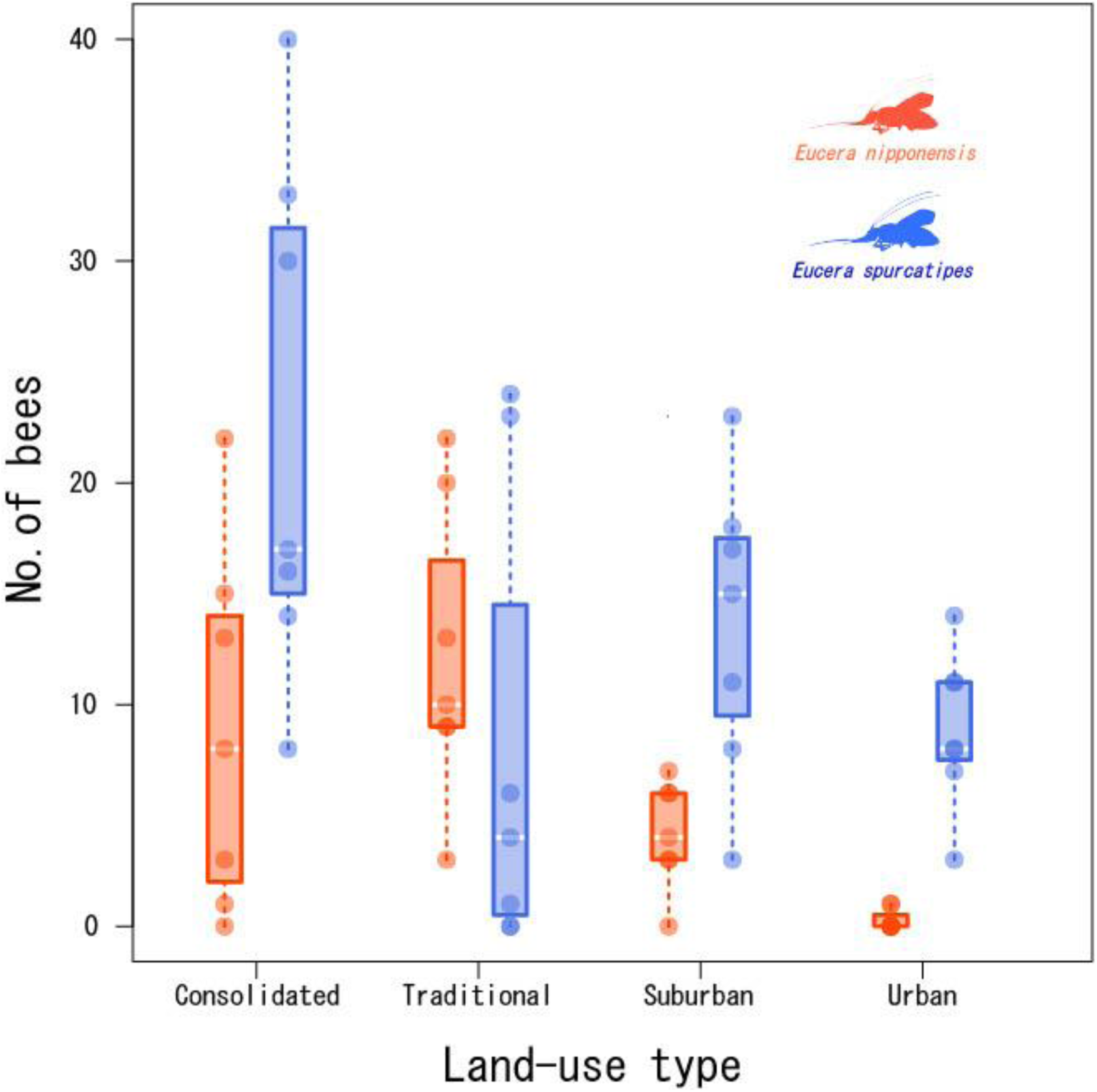
Abundances of the two long-horned bees in paddy meadow study sites with different land uses (orange circles, *Eucera nipponensis*; blue circles, *E. spurcatipes*). Box plots represent the median (bold white horizontal line), first and third quartiles represent box perimeters, and the whiskers represent 90th percentiles.

**Fig. 2.**
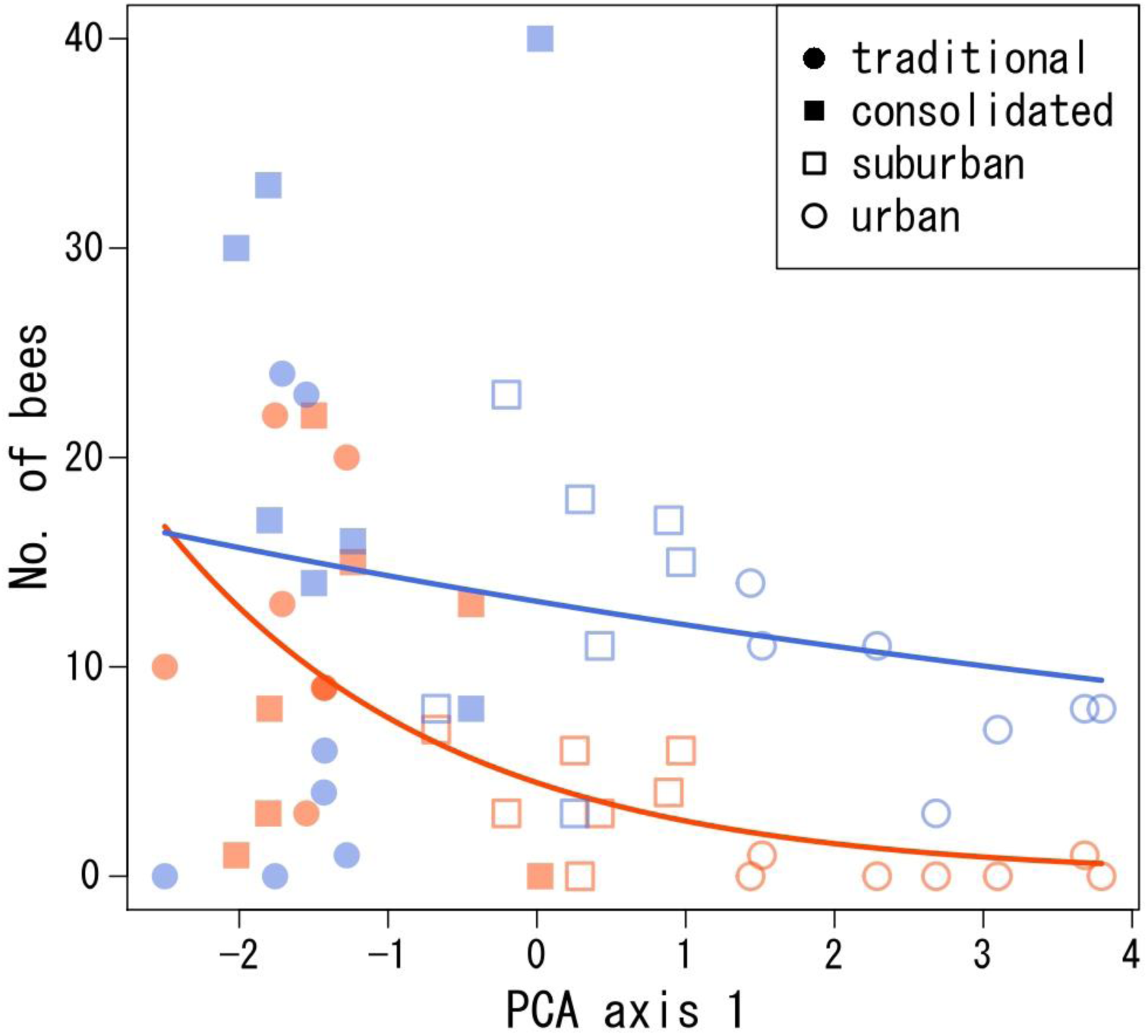
Relationships between the two long-horned bee abundances (orange, *Eucera nipponensis*; blue, *E. spurcatipes*) and the PC1 value in 28 study sites: traditional (closed circle), consolidated (closed square), suburban (open square), and urban sites (open circle). The solid lines (orange, *E. nipponensis*; blue, *E. spurcatipes*) represent significant regressions estimated from the GLM (Table S1).

**Fig. 3.**
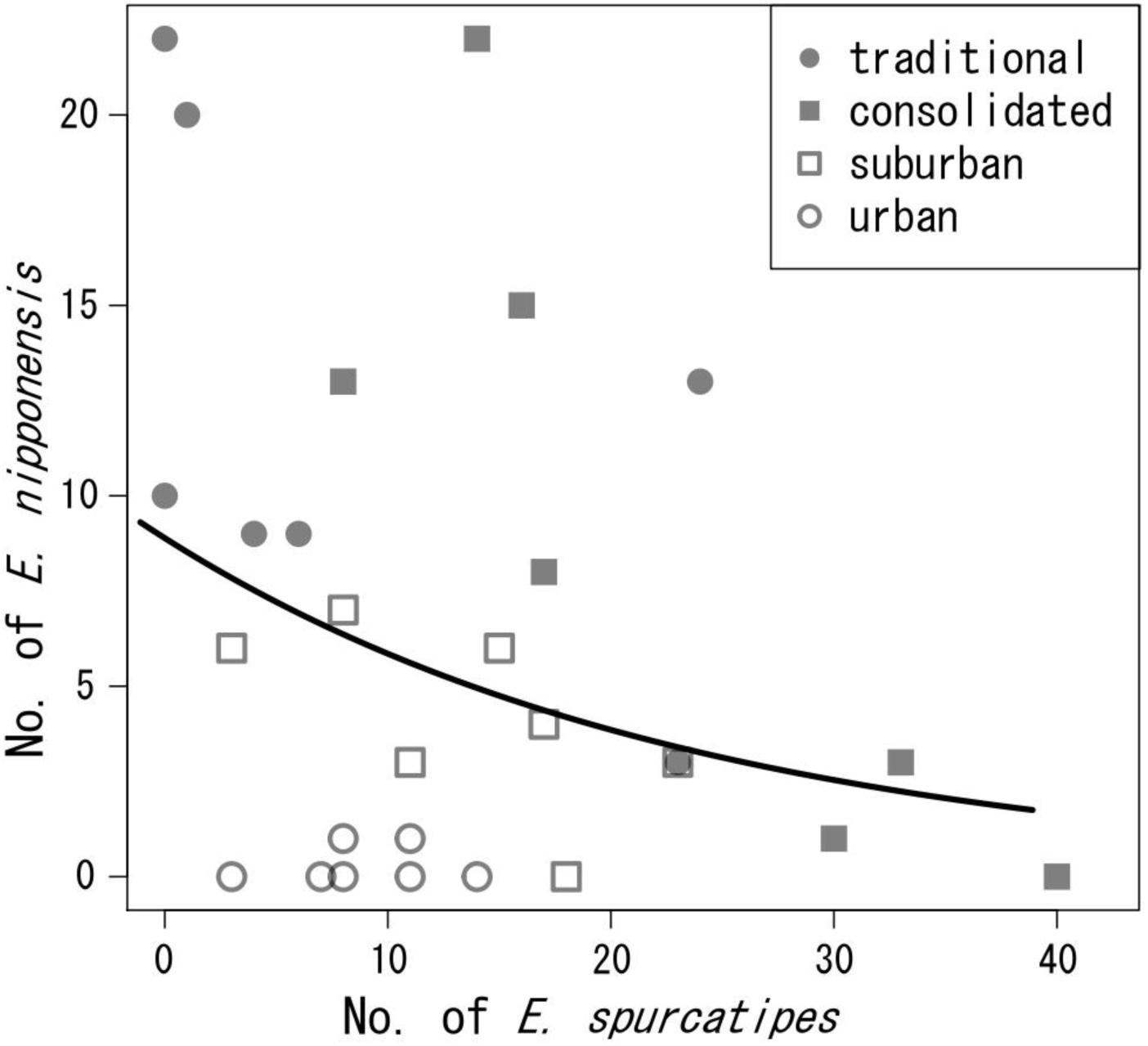
Negative relationship between the abundances of *Eucera nipponensis* and *E. spurcatipes* at 28 study sites: traditional (closed circle), consolidated (closed square), suburban (open square), and urban (open circle). The line represents a significant regression estimated from the GLM (Table S1).

### Flower use patterns

In total, we recorded 61 flowering plant species and observed 267 visits by the two *Eucera* species to 16 species during the study. Flower use patterns significantly differed between the two *Eucera* species in the traditional and consolidated sites (Fig. 4; Fisher’s exact test, *p<0*.*001* for each type). Specifically, *E. nipponensis* visited a variety of flowering species in traditional sites, whereas *E. spurcatipes* used only a limited number of plant species (Fig. 4). Overall, native *Vicia sativa L*. and exotic *Trifolium repens L*. were the most frequently visited flowering species in this study (86 and 110 visits, respectively).

**Fig. 4.**
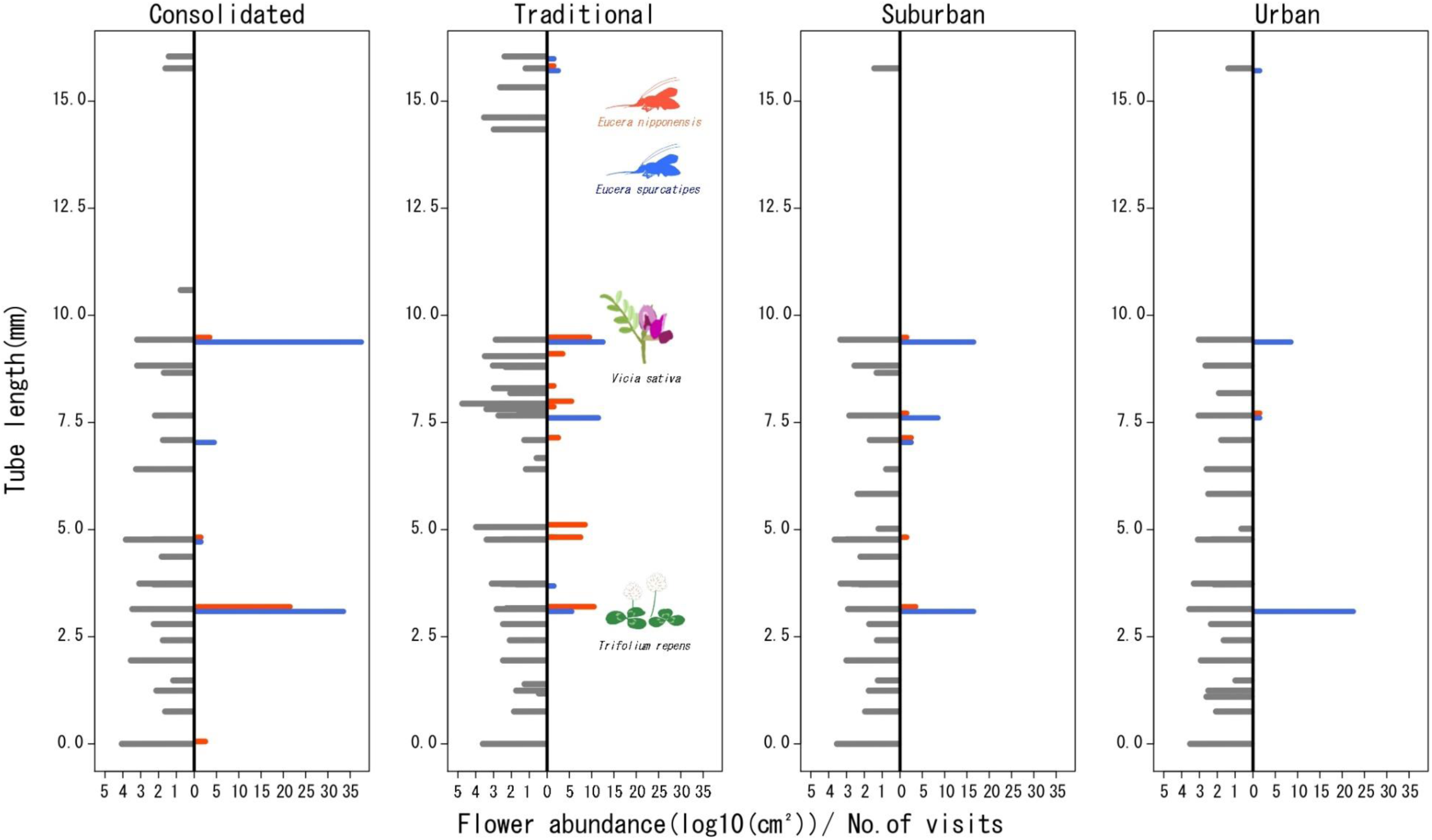
Flower use patterns of *Eucera nipponensis* and *E. spurcatipes* in four different land-use types. The left grey bars of each plot show background flower abundances of species with different tube lengths, which was calculated as the total visual size of flowers (the mean visual size multiplied by the number of flowers). The right orange and blue bars show visit frequencies of *E. nipponensis* and *E. spurcatipes* to each flower species, respectively. Native (*Vicia sativa*) and exotic (*Trifolium repens*) legumes were the most visited flower species in the study.

## Discussion

Our results support the prediction that interspecific differences with regard to the susceptibility of a species to land-use changes are influenced by habitat preference differences. We found that *E. nipponensis* was less likely to occur in consolidated and urbanised meadows, whereas *E. spurcatipes* maintained and increased its abundance in these meadows. We also found that the different response of land-use change likely caused changes of surrounding landscape for urbanisation and floral resources for paddy consolidation. In the following discussion, we consider the effects of land-use changes on pollinators with different habitat and flower preferences.

The effects of land-use changes on pollinators can vary with the type of habitat transitioned to and from, depending on species characteristics (Winfree et al. 2011). Our results showed similar but slightly different effects of urban landscapes on both *Eucera* species; forest-associated *E. nipponensis* was more vulnerable to decrease and increase in surrounding forests and developed lands, respectively, than open land-associated *E. spurcatipes* (Fig 2). Moreover, *E. spurcatipes* became more abundant in both consolidated and urbanised sites than *E. nipponensis* (Fig. 1), indicating that current intensive land-use changes positively affected the relative dominance of *E. spurcatipes*. Ground-nesting solitary bees are usually more susceptible to changes in land use and management associated with agricultural intensification (Williams et al. 2010). Our findings partly support the previous study and suggest that solitary bees can adapt to intensified agricultural lands when they prefer disturbed habitats. Such species-specific responses to land-use changes often cause compositional shifts in pollinator communities. In our case, intensive land-use changes would promote the replacement of forest-associated species by open land-associated species.

Floral resources are also an important determining factor of pollinator distributions (Roulston and Goodell 2011), and land-use changes often alter flowering resource composition, which in turn can affect pollinator distribution (Winfree et al. 2011). In our study, *E. nipponensis* and *E. spurcatipes* showed a strong preference for legume flowers (63.5% and 92.9%, respectively), such as *V. sativa* and *T. repens*, which were abundant in all types of paddy fields (Fig. 4). The distribution and abundance of the study bees may have been unaffected by floral resource composition in this study. However, *E. nipponensis* visited a variety of flowers in traditional sites and exhibited a wider diet breadth than *E. spurcatipes* (Fig. 4). Specialist pollinators with narrow diet breadth are usually more vulnerable to land-use change than generalists because of the loss of their main host plants (Winfree et al. 2011; Weiner et al. 2014). Contrary to this common assumption, legume-specialised *E. spurcatipes* were more robust to land-use changes than generalist *E. nipponensis* in our study. The consistent presence of legume flowers at the study sites could have supported populations of *E. spurcatipes*. In contrast, poor flower richness in consolidated and urbanised paddy meadows (Uchida & Ushimaru 2014; Uchida et al. 2018; this study) likely reduced the diet breadth of *E. nipponensis* and potentially caused population shrinkage in this species.

We found a negative relationship between *E. nipponensis* and *E. spurcatipes* abundance in the study area (Fig. 3), indicating that interspecific competition existed between the species. Interspecific competition for floral resources usually occurs when pollinators’ foraging preferences overlap (Stout and Morales 2009), but the competition would be relaxed when the species forage from a broader range of flowers (Goulson et al. 2008). In our study, *E. nipponensis* had a broader foraging range and different flower use patterns in consolidated and traditional meadows compared to *E. spurcatipes* (Fig. 5). In areas where both bee species coexisted in abundance, *E. nipponensis* frequently visited woody plants and shallow dish-shaped flowers in addition to legume flowers (GSH, personal observation). Similar flower use patterns of both *Eucera* species were recorded in kaki persimmon orchards (Nikkeshi et al. 2021). Thus, our results do not support the existence of strong competition for floral resources between both species. The strength of interspecific competition among bees may vary depending on floral availability and the flexibility of pollinator foraging (Shibata and Kudo 2020). Thus, flexible flower use by *E. nipponensis* would have reduced competition with *E. spurcatipes*.

This study provides a first step in addressing the impact of multiple land-use changes on pollinators in paddy landscapes. We quantitatively revealed that forest-associated species have declined owing to the intensified use of surrounding landscapes in consolidated and urbanised paddy fields, while a species that prefers disturbed habitats has increased in population size. Intensified and urbanised agricultural lands could be alternative habitats for some pollinators (Goddard et al. 2010; Burkle et al. 2017), however, these areas may not be suitable habitats for forest bee community (Harrison et al. 2018). Thus, the recent global prevalence of agricultural intensification and urbanisation could have different impacts on species with different habitat preferences and primarily decrease forest-associated species. To validate the generality of our findings, a comparison of more closely related species preferring different habitats is required worldwide.

## Acknowledgements

We are grateful to Prof. Hiroshi S. Ishii (Toyama University) for his suggestions on bee capturing method. We also thank members of the Biodiversity Laboratory at Kobe University for their valuable comments on our study.

