## Supplemental materials for "Habitat preference influences response to changing agricultural landscapes in two long-horned bees"

**Supplemental material**

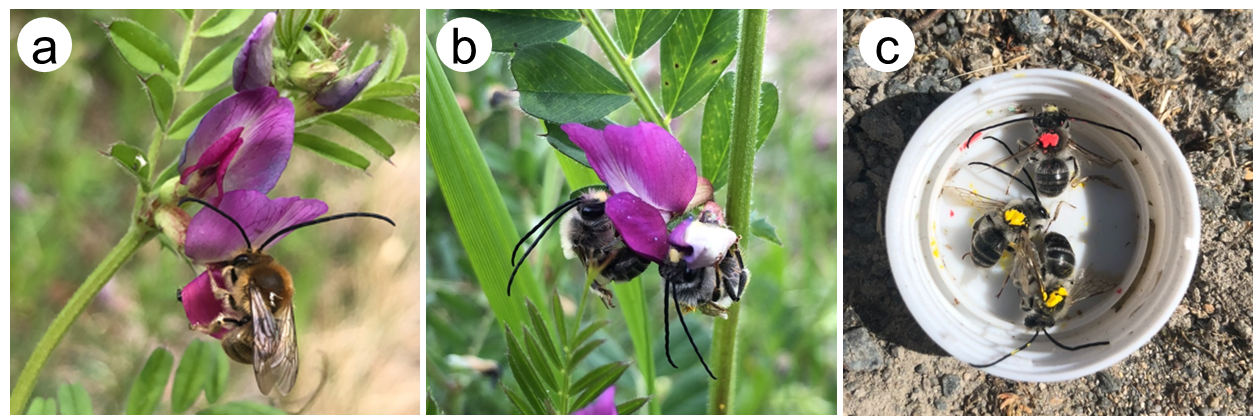

**Fig. S1** Photos of study species: (a) *Eucera nipponensis* visiting *Vicia sativa*; (b) *E. spurcatipes* visiting *V. sativa*; (c) *Eucera* species marked on their thorax by permanent marker (red, *E. nipponensis*; yellow, *E. spurcatipes*)

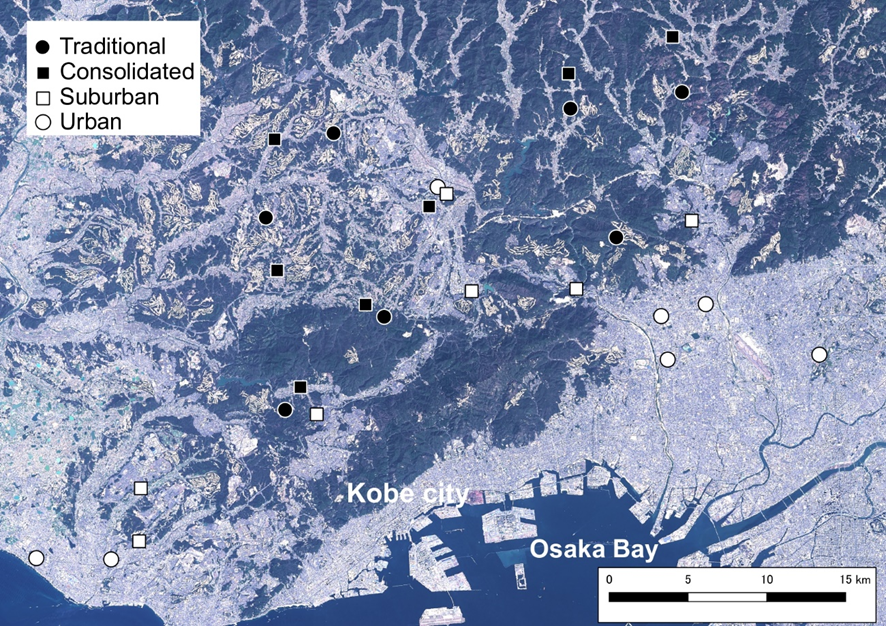

**Fig. S2** Map of the 28 paddy study sites in the Osaka-Kobe metropolitan area, Japan. Seven traditional (closed circle), seven consolidated (closed square), seven suburban (open square), and seven urban (open circle) sites were surveyed.

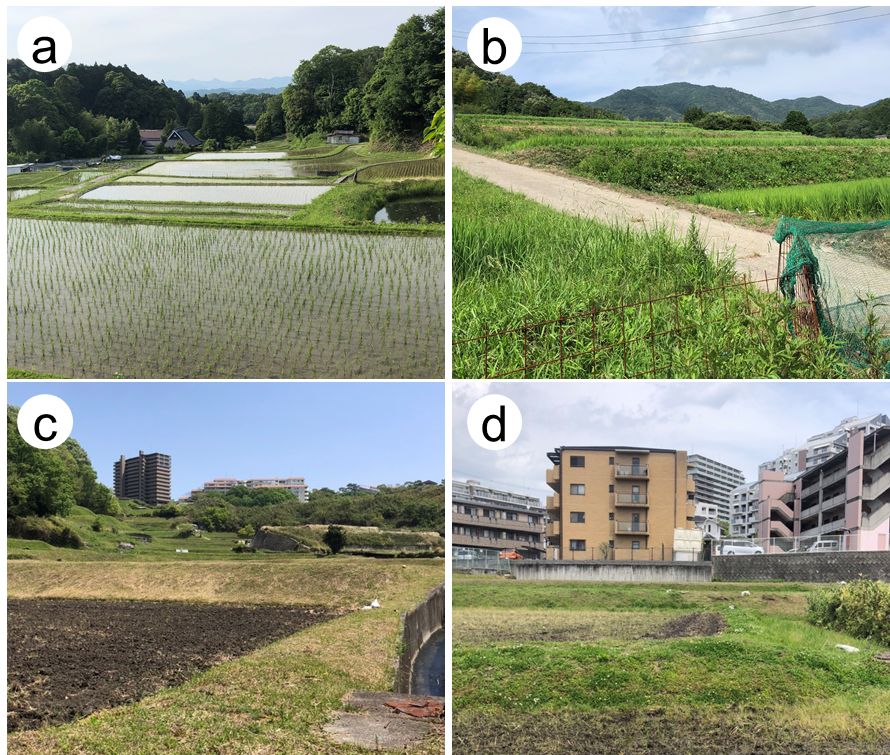

**Fig.S3** Photos of four land-use type of study sites: (a) Traditional (b) Consolidated (c) Suburban (d) Urban.

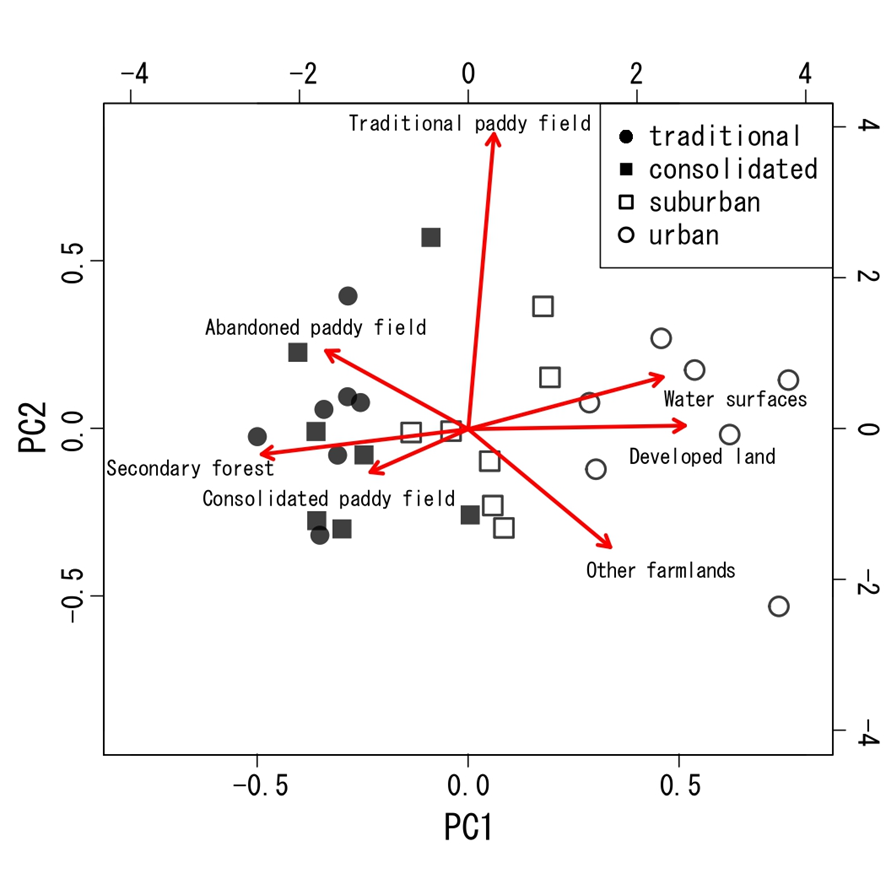

**Fig. S4** PCA bi-plot for landscape variables (calculated from areas of developed lands; secondary forests; traditional, consolidated, and abandoned paddy fields; other farmlands; and water surfaces). The first two PCA axes explain 63.2% of the total variance (PC1: 46.8%, PC2: 16.4%). PC1 was positively correlated with developed land, water surfaces, and other farmlands and negatively correlated with secondary forests and consolidated and abandoned paddy fields. Meanwhile, PC2 was positively correlated with traditional paddy fields and water surfaces and negatively with other farmland areas.

**TableS1** Results of generalized linear model (GLM) analysis. Traditional meadows and *E. nipponensis* were used as baselines for the land-use type and species identity variables, respectively in the models. Significant effects are indicated in bold (*P* < 0.05).

| Response variable | Explanatory variable | Coefficients | | *z* | *P* |
| --- | --- | --- | --- | --- | --- |
|  |  | Estimate | SE |  |  |
| No. of *Eucera* bees | **Land-use type (consolidated)** | -0.334 | 0.169 | -1.971 | **0.0487** |
|  | **Land-use type (suburban)** | -1.096 | 0.218 | -5.022 | **<0.001** |
|  | **Land-use type (urban)** | -3.777 | 0.719 | -5.254 | **<0.001** |
|  | **species identity (*E. spurcatipes*)** | -0.394 | 0.170 | -2.318 | **0.0204** |
|  | **Land-use (consolidated)×species (*E.spurcatipes*)** | 1.329 | 0.227 | 5.868 | **<0.001** |
|  | **Land-use (suburban)×species (*E.spurcatipes*)** | 1.580 | 0.272 | 5.815 | **<0.001** |
|  | **Land-use (urban)×species (*E.spurcatipes*)** | 3.828 | 0.738 | 5.185 | **<0.001** |
|  | Spatial autocovariates | -0.004 | 0.017 | -0.222 | 0.8247 |
|  | **Intercept** | 2.555 | 0.234 | 10.913 | **<0.001** |
| No. of *Eucera* bees | **PC1 values** | -0.527 | 0.066 | -7.945 | **<0.001** |
|  | **species identity (*E. spurcatipes*)** | 1.078 | 0.119 | 9.023 | **<0.001** |
|  | **PC1×species (*E. spurcatipes*)** | 0.438 | 0.073 | 6.028 | **<0.001** |
|  | PC2 values | -0.056 | 0.046 | -1.225 | 0.221 |
|  | Spatial autocovariates | 0.002 | 0.018 | 0.101 | 0.92 |
|  | **Intercept** | 1.479 | 0.210 | 7.032 | **<0.001** |
| No. of *E. nipponensis* | **No. of *E. spurcatipes*** | -0.042 | 0.010 | -4.372 | **<0.001** |
|  | **Spatial autocovariates** | 0.122 | 0.019 | 6.299 | **<0.001** |
|  | **Intercept** | 1.491 | 0.195 | 7.662 | **<0.001** |
